# Automated Calculation of the Disruption Index: A Reproducible Computational Workflow for Large-Scale Bibliometric Analyses

**DOI:** 10.64898/2026.02.12.705484

**Authors:** Analina Braga Apolinário, Kaio Vinicius Vieira, Ana Karolina Mendes Mathias Costa, Larissa Costa Freitas, Isabela Sales Pinheiro, Robert Willer Farinazzo Vitral, Marcio José da Silva Campos

## Abstract

Bibliometric analyses have become essential for understanding scientific production and innovation dynamics; however, large-scale applications remain limited by challenges related to data extraction, preprocessing, citation network reconstruction, and reproducibility, particularly when using PubMed-indexed records. This study presents a fully automated and reproducible computational workflow for large-scale bibliometric analyses based on the Disruption Index (DI). The pipeline enables systematic retrieval of PubMed data, standardized metadata processing, construction of citation networks, and calculation of DI values within a fixed post-publication citation window. Implemented in Python, the workflow integrates automated querying, XML parsing, data consolidation, and network-based citation classification, allowing scalable and transparent analyses that are infeasible through manual approaches. In a demonstrative application focused on orthodontic literature, the pipeline processed more than 67,000 articles and reconstructed over 300,000 citation relationships, resulting in a final analytical sample of 3,234 articles with indexed references and citations. The automated framework ensures methodological transparency, facilitates replication, and substantially reduces the time and technical barriers associated with advanced bibliometric studies. By providing an open and extensible solution for calculating the Disruption Index at scale, this workflow supports robust assessments of scientific innovation and consolidation and can be readily adapted to other biomedical research domains indexed in PubMed.

## INTRODUCTION

Bibliometrics, or the “science of science,” has gained increasing importance by enabling the systematic analysis of scientific dynamics, providing support to understand how academic production functions, to guide science and technology policies, and to assess the social impact of research [1]. The substantial growth in the volume of publications over recent decades has only become analyzable due to advances in large-scale databases and the development of computational tools capable of processing massive information sets [1,3]. In this context, automation has played a decisive role by transforming analyses that were previously infeasible—because of their complexity, time requirements, or high degree of manual intervention—into procedures that are accessible, precise, and reproducible [4,5].

The reliability of any bibliometric analysis depends directly on the database used. Scopus and Web of Science stand out for their broad multidisciplinary coverage, stable performance, and integrated analysis tools [6,7]. However, in the biomedical field, PubMed holds a central position due to the breadth of its collection, open access, and the structured use of MeSH descriptors, which contribute to more refined and consistent searches [7,8]. Despite these advantages, advanced analyses based on raw PubMed data still face significant barriers: heterogeneous metadata, the need for standardization, and preprocessing steps are often not described in sufficient detail in scientific articles, hindering replicability and creating methodological gaps that affect transparency and the scope of studies [1,9].

These limitations become even more evident given that many traditional metrics rely exclusively on citation counts. Although widely used, citations alone are insufficient to reveal why a work is cited and, therefore, cannot capture nuances related to scientific innovation [1,2,10]. The literature highlights that highly innovative articles play a critical role in scientific progress and may trigger paradigm shifts, yet they are often not identified by traditional indicators [11,12]. With the aim of quantifying this innovative behavior—or, conversely, consolidative behavior—Funk and Owen-Smith (2016) proposed the Disruption Index (DI) [13].

The DI simultaneously incorporates relationships between an article, its references, and the citations it receives, allowing one to distinguish works that redirect attention toward new research trajectories (disruptive) from those that reinforce established paths (consolidative) [12–14]. Unlike metrics based solely on the total number of citations, the DI considers co-citation patterns among subsequent works, offering a more robust measure of a publication’s structural influence on the development of science [10,14]. The indicator ranges from –1 to +1, reflecting consolidative and disruptive behaviors, respectively, and has been applied to study phenomena from technological change to the dynamics of scientific teams [12,15].

Although conceptually powerful, large-scale DI computation still faces practical challenges. The process requires building extensive citation networks, accurately identifying relationships between later articles and their references, and defining appropriate time windows—often six years— considered sufficient for citation stabilization in health-related fields [14,16]. In addition, standardized, automated, and reproducible pipelines that enable consistent application of the DI across large biomedical databases remain scarce, which limits its systematic adoption.

Against this backdrop, the present study aimed to develop a fully automated and reproducible computational workflow to extract PubMed data, build citation networks, and calculate the Disruption Index at scale. The system was implemented in Python, supported by artificial intelligence tools, and made openly available to ensure transparency and replicability [17]. Although demonstrated through an applied study, the pipeline was designed to be universal and adaptable to different scientific domains.

## METHODOLOGY

### 2.1. Study Design

This study aimed to develop and validate a fully automated computational workflow for PubMed data extraction, metadata organization, citation-network reconstruction, and Disruption Index (DI) computation, in line with the literature on scientific innovation metrics [1–3,12–15]. The entire pipeline was implemented in Python, executed in a Jupyter Notebook environment, and supported by automation of critical steps to ensure reproducibility and minimize manual intervention.

Although the illustrative application focuses on publications related to orthodontics, the pipeline was designed to be methodologically universal and can be applied to any biomedical domain indexed in PubMed.

### 2.2. Search Strategy and Publication Selection

Publications were identified in PubMed (https://pubmed.ncbi.nlm.nih.gov/) through an automated search using the term “Orthodontics” applied to the title, abstract, or MeSH descriptors. After retrieval, only articles that met all of the following criteria were included:

- had references indexed in PubMed;
- had citations also indexed in PubMed;
- allowed reconstruction of citation relationships;
- had a citation window compatible with the method.

Articles lacking references, lacking citations, or presenting insufficient metadata were automatically excluded by the preprocessing script. In accordance with the literature, a fixed six-year window was used for counting subsequent citations.

### 2.3. Collection of Identifiers (PMIDs)

The collection of unique article identifiers (PMIDs) was carried out automatically using the NCBI E-utilities API. The script employed (Appendix A — Supporting Information) performs:

- automated querying of PubMed;
- pagination of results;
- extraction of PMIDs;
- classification of records by year;
- saving identifiers to .csv files.

This process establishes the initial document base for all subsequent steps of the workflow.

### 2.4. Extraction of XML Files and Metadata

For each retrieved PMID, complete metadata were extracted using the E-fetch service. The corresponding script (Appendix B) performs:

- downloading of XML files;
- identification and extraction of relevant fields;
- organization of files into folders structured by year.

The extracted metadata include:

- title;
- year and date of publication;
- journal and ISSN;
- authors and affiliations;
- ORCID (when available);
- MeSH descriptors;
- publication type;
- language;
- country;
- funding information;
- editorial dates (submission, acceptance, and publication);
- cited references (PMIDs);
- PubMed-indexed citations.

These elements constitute the raw dataset for standardization and analysis.

### 2.5. Data Cleaning, Standardization, and Consolidation

Metadata consolidation was performed using a dedicated script (Appendix C), responsible for:

- reading and parsing XML files;
- transforming metadata into a standardized dataframe;
- harmonizing redundant fields;
- structuring all data into a consolidated spreadsheet.

This spreadsheet includes:

- PMID;
- year;
- title;
- list of authors and affiliations;
- ORCID;
- MeSH descriptors;
- journal and ISSN;
- publication type;
- country and language;
- funding agencies;
- complete list of references (cited PMIDs).

### 2.6. Construction of the Citation Network

The citation network was built as described in Appendix D. For each article, the algorithm identifies:

- received citations;
- all cited references;
- cross-links between later articles and the references of each focal article.

This reconstruction enables the definition of three sets essential to DI computation:

- A — articles that cite exclusively the focal article;
- B — articles that cite the focal article and at least one of its references;
- C — articles that cite only the focal article’s references.

Networks were constructed respecting a six-year window after publication of the focal article, as established in the literature [12–16].

### 2.7. Disruption Index Calculation

The disruption_index function used in this stage is derived from the *pySciSci* package, as described by Gates and Barabási [18]. The equation employed is grounded in Wu, Wang, and Evans (2019) [15].

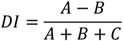

Where:

- A = citations exclusively to the focal article;
- B = citations jointly to the focal article and to at least one of its references;
- C = citations made exclusively to the focal article’s references.

The limits and general properties of the DI were described by Funk and Owen-Smith (2016) [13], Sullivan et al. (2021) [2], and Leibel & Bornmann (2024) [14].

The script:

- counts A, B, and C;
- applies the temporal window;
- checks the consistency of citation relationships;
- generates a final spreadsheet containing:
  - PMID,
  - year,
  - A, B, C,
  - total citations,
  - DI value.

### 2.8. Pipeline Validation

Validation was conducted through:

1. Verification of citation counts A set of randomly selected articles had their citation counts manually compared with those displayed in PubMed. The correspondence confirmed the fidelity of the network reconstruction.
2. Manual verification of the equation The DI equation was solved manually for selected articles. The manually derived values matched the script outputs, confirming the accuracy of the automated calculation.

### 2.9. Reproducibility and Availability

All scripts used—search, extraction, standardization, analysis, and DI calculation—are available in *SciDisruptor-PubMed* [17], ensuring full reproducibility of the pipeline.

## RESULTS

The computational pipeline developed in this study, illustrated in Figure 2, comprised five sequential stages. The initial stage, used for demonstrative purposes, identified 67,632 PMIDs associated with the term “Orthodontic.” These records subsequently underwent the retrieval and cleaning procedures corresponding to the second and third stages of the computational workflow.

Citation-network construction, performed in the fourth stage, applied a fixed six-year window after publication (Figure 1). The resulting analysis incorporated 303,700 citations, enabling reconstruction of relationships between citing articles and referenced works. After integrating metadata with the citation network, 3,234 articles were identified as having both references and citations indexed in PubMed, forming the final sample used for Disruption Index calculation.

**Figure 1.**
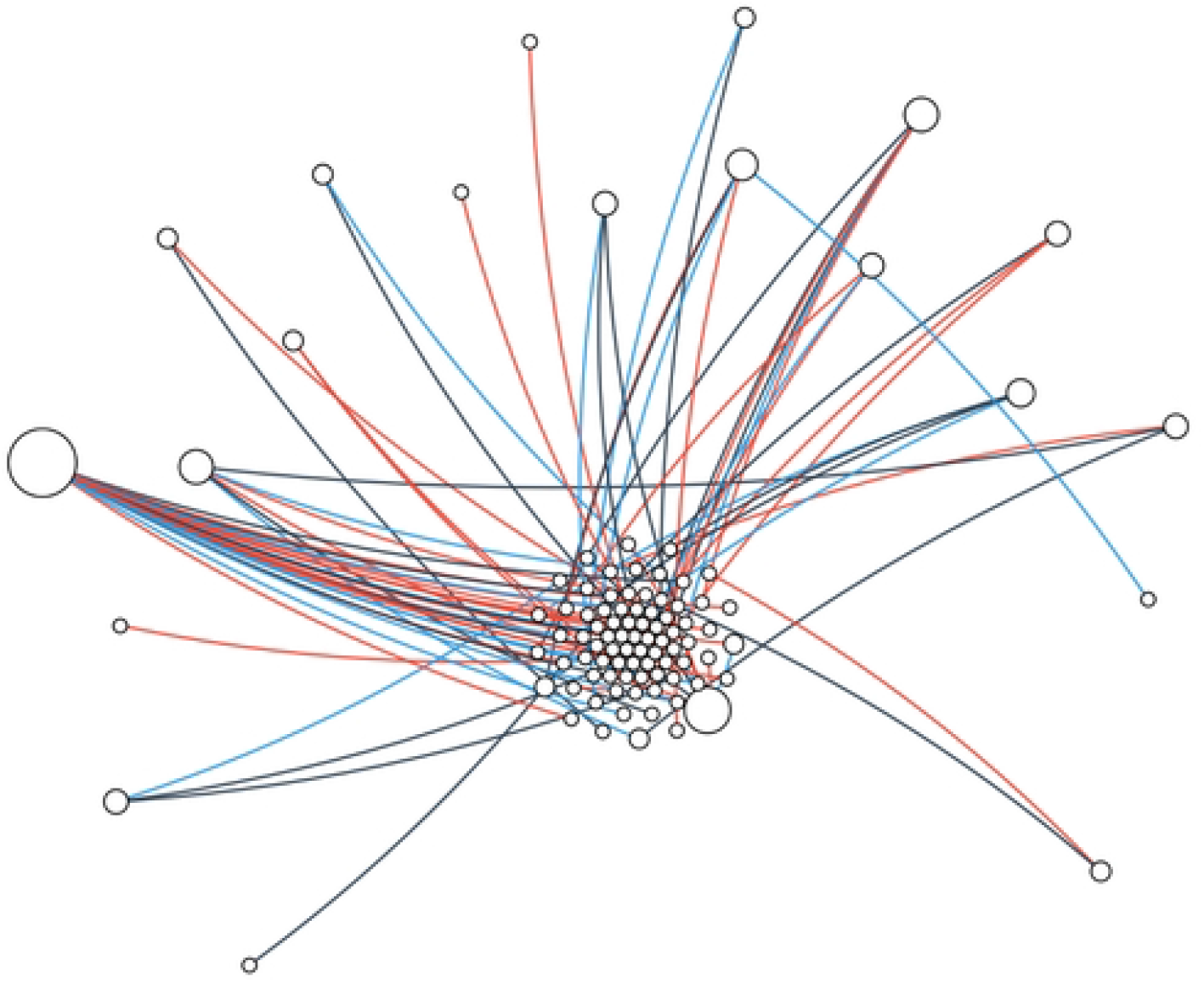
Schematic representation of the citation network used for Disruption Index classification. This figure illustrates a schematic citation network constructed for the calculation of the Disruption Index (DI). The central node represents a focal article, while surrounding nodes correspond to subsequent articles that cite the focal article and/or its references. Edges represent citation relationships and are classified according to citation type: Type A citations (blue edges) indicate articles that cite only the focal article; Type B citations (red edges) represent articles that cite both the focal article and at least one of its references; and Type C citations (black edges) correspond to articles that cite only the references of the focal article. Node sizes and spatial arrangement are illustrative and do not represent quantitative citation counts or temporal ordering. The network is shown for conceptual purposes to demonstrate the logic underlying DI computation.

In the final stage, the pipeline generated a .csv output file containing the Disruption Index (DI) values for each of the 3,234 articles, along with the specific counts of citation types (A, B, and C).

**Table 1.** Excerpt from the dataset showing PMIDs, publication years, Type A/B/C citation counts, total citations, and Disruption Index (DI) values.

| PMID | Year | A | B | C | Total Citations | DI |
| --- | --- | --- | --- | --- | --- | --- |
| 30174498 | 2018 | 20 | 0 | 6 | 26 | 0.77 |
| 30173318 | 2019 | 2 | 0 | 7 | 9 | 0.22 |
| 30171391 | 2018 | 11 | 2 | 114 | 127 | 0.07 |
| 30171364 | 2018 | 3 | 0 | 20 | 23 | 0.13 |
| 30167795 | 2019 | 3 | 0 | 16 | 19 | 0.16 |
| 30151384 | 2018 | 2 | 0 | 1 | 3 | 0.67 |
| 30147398 | 2018 | 5 | 10 | 76 | 91 | -0.05 |

## DISCUSSION

The findings of this study show a pattern consistent with the literature on the growth of global scientific output [15,12], including in the health sciences [19,20] and, more specifically, in orthodontics. In the dataset analyzed, a progressive increase in publication volume was observed up to 2015, followed by a decline—an outcome compatible with previous analyses of specialized dental journals [21].

The increasing use of bibliometric analyses has been reported in the literature [2]. The validity of such analyses depends on the availability of large volumes of data, often accessed through computational tools [3], automated systems [5], and advanced algorithms [4]. Web of Science and Scopus are widely used [6,7]; however, in biomedical research, PubMed is considered one of the most suitable databases [8,7,1].

Despite the popularity of traditional metrics such as the h-index [2], citation counts do not distinguish true impact [1] nor do they capture structural innovation. Innovative publications play a critical role in scientific advancement [10,2], and innovation is recognized as a key driver of progress [11]. In this context, the Disruption Index has been applied to study scientific innovation [13,15].

The DI was initially developed for patents [13] and later adapted for scientific publications [15], proving useful for capturing the opening of new research trajectories and the consolidation of knowledge [14]. Interdisciplinary comparisons of DI values are generally discouraged [14,22].

In the sample analyzed, 64.4% of articles exhibited positive DI values, indicating a disruptive nature. Both forms of scientific production—consolidative and disruptive—are important for the progress of science [13,15,12].

The quality of DI estimation depends on the coverage and accuracy of the underlying data [13,10,14]. Limited coverage can introduce distortions, artificially inflating DI values [22], a phenomenon observed in decades prior to 1990. After that period, the increased availability of data [13,10,14] led to a higher number of articles for which DI could be computed. After 2009, a stabilization was observed, with mean DI values ranging approximately between +0.10 and +0.25.

## CONCLUSION

The computational workflow presented in this study provides a robust, transparent, and fully automated methodological framework for extracting PubMed data, reconstructing citation networks, and calculating the Disruption Index at scale. The pipeline’s ability to efficiently process extensive datasets — more than 67.000 articles and over 300.000 citation relationships — highlights its practical relevance and underscores the infeasibility of manual approaches for analyses of this magnitude.

By publicly releasing all scripts and documenting each step in detail, this work directly contributes to strengthening reproducibility in the science of science. The pipeline not only reduces the time and effort required to organize complex metadata but also promotes methodological standardization that is essential for advanced bibliometric analyses, particularly in biomedical contexts.

The resulting framework enables further investigations into disruption patterns across scientific domains, thematic trends, editorial behaviors, and the evolution of scientific output. By incorporating the Disruption Index as a central metric, the pipeline offers a tool capable of identifying both innovative contributions and consolidative trajectories, providing relevant support for strategic analyses, editorial decision-making, and the development of funding policies.

Overall, the results demonstrate that integrating automation, large-scale data, and innovation-oriented metrics represents a promising path to improve the precision, scalability, and replicability of contemporary bibliometric analyses. This approach helps consolidate the Disruption Index as a valid instrument for understanding scientific innovation dynamics and reinforces the importance of open, accessible computational tools for advancing research in the science of science.

## DECLARATIONS

### Funding

This study did not receive any specific funding for research activities or publication costs. The authors acknowledge the Coordination for the Improvement of Higher Education Personnel (CAPES, Finance Code 001) and the Minas Gerais State Agency for Research and Development (FAPEMIG) for providing institutional graduate scholarships, which supported academic training but did not fund this research or its publication.

### Data Availability Statement

All scripts, datasets, and documentation used in this study are publicly available. The dataset and associated Python scripts for data retrieval, processing, citation network construction, and Disruption Index calculation are available at:

SciDisruptor-PubMed [dataset]. Mendeley Data, V1.

https://doi.org/10.17632/hyg4vfgjtn.1

### Ethics Statement

Ethical approval was not required for this study, as it was based exclusively on publicly available bibliographic data and did not involve human participants, animals, or identifiable personal information.

### Competing Interests

The authors declare no competing interests.

### Declaration of Generative AI and AI-Assisted Technologies

During the preparation of this manuscript, the authors used ChatGPT (OpenAI) to assist with language editing and technical writing. The authors reviewed and edited the content as necessary and take full responsibility for the content of the published article.

## Notes

### Competing Interest Statement

The authors have declared no competing interest.

## REFERENCES

1- Liu L, Jones BF, Uzzi B, Wang D. Data, measurement and empirical methods in the science of science. Nature Human Behaviour. 2023;7:1046–1058. doi:10.1038/s41562-023-01562-4

2- Sullivan GA, Skertich NJ, Gulack BC, Becerra AZ, Shah AN. Shifting paradigms: The top 100 most disruptive papers in core pediatric surgery journals. Journal of Pediatric Surgery. 2021;56(8):1263–1274. doi:10.1016/j.jpedsurg.2021.02.00

3- Santo Fortunato et al. Science of science. Science. 2018;359:eaao0185. doi:10.1126/science.aao0185

4- Wong KF, Lam XY, Jiang Y, Wang D, Chan T. Artificial intelligence in orthodontics and orthognathic surgery: a bibliometric analysis of the 100 most-cited articles. Head & Face Medicine. 2023;19:38. doi:10.1186/s13005-023-00383-0

5- Cooper ID. Bibliometrics basics. Journal of the Medical Library Association. 2015;103(4):217–218. doi:10.3163/1536-5050.103.4.013

6- Pranckutė R. Web of Science (WoS) and Scopus: the titans of bibliographic information in today’s academic world. Publications. 2021;9:12. doi:10.3390/publications9010012

7- AlRyalat SAS, Malkawi LW, Momani SM. Comparing bibliometric analysis using PubMed, Scopus, and Web of Science databases. Journal of Visualized Experiments. 2019;(152). doi:10.3791/58494

8- Falagas ME, Pitsouni EI, Malietzis GA, Pappas G. Comparison of PubMed, Scopus, Web of Science, and Google Scholar: strengths and weaknesses. FASEB Journal. 2008;22(2):338–342. doi:10.1096/fj.07-9492LSF

9- Lin Z, Yin Y, Liu L, Wang D. SciSciNet: A large-scale open data lake for the science of science research. Scientific Data. 2023;10:315. doi:10.1038/s41597-023-02198-9

10- Bornmann L, Devarakonda S, Tekles A, Chacko G. Are disruption index indicators convergently valid? Quantitative Science Studies. 2020;1:1242–1259.

11- Hofstra B, et al. The diversity–innovation paradox in science. Proceedings of the National Academy of Sciences. 2020;117:9284–9291.

12- Park M, Leahey E, Funk RJ. Papers and patents are becoming less disruptive over time. Nature. 2023;613:138–144. doi:10.1038/s41586-022-05543-x

13- Funk RJ, Owen-Smith J. A dynamic network measure of technological change. Management Science. 2016;63(3):791–817.

14- Leibel C, Bornmann L. What do we know about the disruption index in scientometrics? Scientometrics. 2024;129:601–639. doi:10.1007/s11192-023-04873-5

15- Wu L, Wang D, Evans JA. Large teams develop and small teams disrupt science and technology. Nature. 2019;566:378–382.

16- Antunes AA. Como avaliar a produção científica. Rev Col Bras Cir. 2015;42(Suppl 1):17–19. doi:10.1590/0100-69912015S01006

17- Apolinário AB, Alves KVV, Costa AKMM, Campos MJS. SciDisruptor-PubMed [dataset]. Mendeley Data. 2025;V1. doi:10.17632/hyg4vfgjtn.1

18- Gates AJ, Barabási A-L. Reproduced science from science at scale: pySciSci. Quantitative Science Studies. 2023;4(3):700–710. doi:10.1162/qss_a_00260

19- Asiri FY, Kruger E, Tennant M. Global dental publications in PubMed databases between 2009 and 2019—A bibliometric analysis. Molecules. 2020;25(20):4747. doi:10.3390/molecules25204747

20- Ali S et al. A bibliometric analysis of scientific publication on peri-implantitis from 1990 to 2020. Journal of Clinical and Experimental Dentistry. 2024;16(5):e570–e579.

21- Almotairy N. International trends of orthodontic publications. Dental Press Journal of Orthodontics. 2023;28(1):e2321175. doi:10.1590/2177-6709.28.1.e2321175.oar

22- Ruan X et al. Rethinking the disruption index as a measure of scientific and technological advances. Technological Forecasting and Social Change. 2021;172:121071. doi:10.1016/j.techfore.2021.121071

